# Oligonucleotides can act as superscaffolds that enhance liquid-liquid phase separation of intracellular mixtures

**DOI:** 10.1101/2020.01.24.916858

**Authors:** Jerelle A. Joseph, Jorge R. Espinosa, Ignacio Sanchez-Burgos, Adiran Garaizar, Daan Frenkel, Rosana Collepardo-Guevara

**Affiliations:** Maxwell Centre, Cavendish Laboratory, Department of Physics, University of Cambridge, J J Thomson Avenue, Cambridge CB3 0HE; Department of Chemistry, University of Cambridge, Lensfield Road, Cambridge, CB2 1EW; Department of Genetics, University of Cambridge, Cambridge, CB2 3EH

## Abstract

Intracellular liquid-liquid phase separation (LLPS) enables the formation of biomolecular condensates, which play a crucial role in the spatiotemporal organisation of biomolecules (proteins, oligonucleotides). While LLPS of biopolymers has been demonstrated in both experiments and computer simulations, the physical determinants governing phase separation of protein-oligonucleotide systems are not fully understood. Here, we introduce a minimal coarse-grained model to investigate concentration-dependent features of protein-oligonucleotide mixtures. We demonstrate that adding oligonucleotides to biomolecular condensates composed of oligonucleotide-binding scaffold proteins enhances LLPS; since oligonucleotides act as ultra-high-valency molecules (termed ‘superscaffolds’) that increase the molecular connectivity among scaffold proteins. Importantly, we find that oligonucleotides promote protein LLPS via a seeding-type mechanism; recruiting numerous protein molecules and reducing the thermodynamic and kinetic barriers for nucleation and phase separation. By probing the conformational properties of oligonucleotides within droplets, we show that these biopolymers can undergo phase separation-driven compaction, which may be entropic in nature. Finally, we provide a quantitative comparison between mixture composition, protein valency, and protein-oligonucleotide interaction strengths. We find that superscaffolds preferentially recruit higher valency proteins to condensates, and that multiphase immiscibility within condensates can be achieved by modulating the relative protein-oligonucleotide binding strengths. These results shed light on the roles of oligonucleotides in ribonucleoprotein granule formation, heterochromatin compaction, and internal structuring of the nucleolus and stress granules.

## Introduction

In recent years, it has become clear that liquid-liquid phase separation (LLPS) is responsible for the formation of membraneless organelles, including P granules, nucleoli, and nuclear bodies (NBs).^1–4^ These cellular bodies, often referred to as biomolecular condensates, display liquid-like properties, such as the ability to flow, coalesce, and drip,^5–9^ and are thought to self-assemble via condensation of proteins and other macromolecules in the cytoplasm and nucleoplasm. Specifically, multidomain proteins^5,8,10^ and those comprising intrinsically disordered regions (IDRs)^11–14^ have been shown to undergo LLPS, both in vitro and in cells. LLPS is mainly driven by multivalent protein-protein interactions. Interactions with oligonucleotides (i.e. RNA and DNA)^15–19^ may also mediate LLPS; indeed, many membraneless organelles are ribonucleoprotein (RNP) granules, consisting of RNA and RNA-binding proteins.^20^ However, the mechanism by which these biopolymers enhance (or inhibit) LLPS is not fully understood. Moreover, the predictive rules governing the composition of protein-RNA/DNA condensates are unclear. There is, therefore, a need for biophysical models that can predict intracellular LLPS and delineate the underlying molecular mechanism; this is a central goal of the present work.

Experiments reveal that, although condensates are multicomponent systems, only a small subset of components may be required for LLPS.^21,22^ For example, P granules are made up of several RNA and protein molecules; however, LAF-1 (a protein found in P granules) can self-associate into P granule-like droplets in vitro.^13^ In some cases, macromolecules that undergo LLPS may bind to and recruit other molecules to phase-separated droplets. Banani et al.^23^ showed that polySUMO and polySIM proteins assembled into droplets when mixed, and subsequently recruited fluorescently labelled SIM and SUMO monomers, respectively, to the condensates. Components, such as polySUMO and polySIM, that drive LLPS are often classified as ‘scaffolds’, and molecules that partition into droplets formed by scaffolds (e.g. SIM and SUMO monomers) are termed ‘clients’.^23^ Scaffold and client stoichiometric ratios, valencies, and binding affinities have been postulated as crucial in compositional control of membraneless organelles.^23^ Simulations can further elucidate the interplay between these factors. We need to understand how oligonucleotides may be selectively recruited to condensates, and how this recruitment is affected by factors such as length, flexibility, and shape.^12,18,24,25^ Furthermore, RNA/DNA themselves may act as scaffolds and seeds for LLPS.^20,26–28^ Thus, oligonucleotides may not just participate in, but even control LLPS.

Polymer physics provides key rules for predicting phase behaviour of polymeric systems.^29^ For a given homopolymer-solvent mixture, the system produces two phases (i.e. a dense, polymer-rich phase and a dilute, solvent-rich phase) when the enthalpy of mixing exceeds the entropy of mixing. The Flory-Huggins Theory quantifies the entropic and enthalpic terms of such systems and estimates the critical condition for phase separation.^30,31^ Other theories have been developed for mixtures containing charged polymers,^32–34^ as well as more generalised models for heteropolymeric systems.^35^ These models can be useful in understanding LLPS of biopolymers. The study of phase diagrams of biopolymer systems can provide valuable insight in the various factors that influence phase separation. However, measuring complete phase diagrams is time consuming and the study of biopolymer phase behaviour in fully atomistic simulations tends to be computationally expensive. Hence, minimalist models that can capture observable phase behaviour in an efficient manner are appealing, as they can provide insight at moderate computational cost.

Here we introduce a simple coarse-grained model for probing phase separation in protein-oligonucleotide mixtures. The model is simple enough to allow simulation of phase transitions in mixtures containing thousands of molecules at low computational cost. We first investigate the effect of adding an oligonucleotide to a pure protein system. We find that the oligonucleotide acts as a “superscaffold”; by enhancing the average valency of the system, it increases the critical temperature for phase separation. These results are consistent with experiments, where RNA was found to decrease the critical concentration for LLPS in mixtures of RNA and RNA-binding proteins.^20,26–28^

Biopolymers recruited to droplets may be more compact than in the cytosol/nucleosol.^24^ An example is the collapsed structure of heterochromatin involved in LLPS.^36^ Although one would expect that polymer compaction decreases the entropy of the system, here we rationalise that the formation of oligonucleotide condensates is possibly entropically driven. Finally, we study competition and cooperative effects in multicomponent mixtures, and demonstrate how the droplet composition is tuned by the valencies, stoichiometries, and relative interaction strengths of the components.

## Results and discussion

### Model

The tendency of a solution to undergo LLPS depends critically on the valencies (number of attractive sites per particle) and relative interaction strengths of the molecular components. Hence, any simple model for LLPS must provide reasonable approximations to these physical and energetic parameters.

In this work, multivalent proteins are modelled as hard spheres decorated with attractive patches or “stickers” (Fig. 1A). Scaffold proteins contain three/four patches (3/4-valency proteins), while, clients are modelled as hard spheres with two attractive sites (2-valency proteins). Oligonucleotides contain negatively charged sugar-phosphate backbones, and therefore tend to behave as self-avoiding polymers. Hence, as an extension to our previous work,^37^ to model oligonucleotides, we introduce flexible, self-avoiding polymers (i.e. chains of hard spheres) that interact with patchy particles via weak attractive interactions (Fig. 1A). The weak attractive interactions between patchy “protein” particles and polymers represent protein-oligonucleotide interactions observed in biomolecular condensates.^20,26–28^

**Figure 1:**
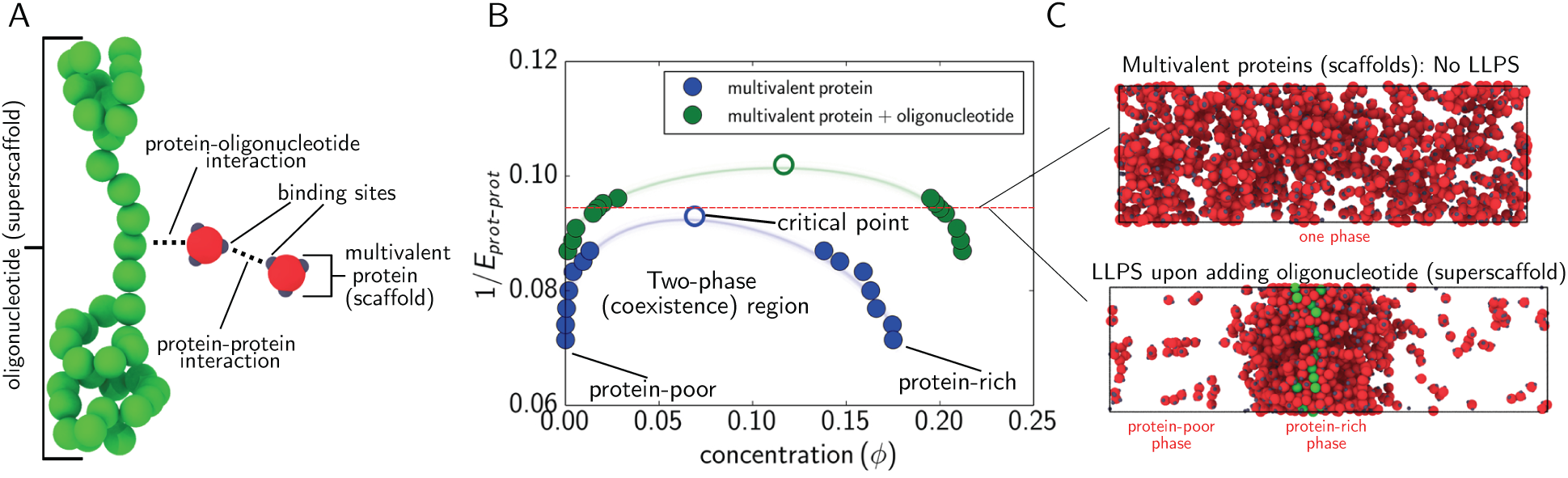
Obtaining phase diagrams of protein-oligonucleotide mixtures. (A) Minimalist coarse-grained model for simulating interactions between proteins and oligonucleotides. (B) Phase diagram, in terms of inverse inter-protein interaction strength (1*/E*_*prot*-*prot*_) and concentration (i.e. packing fraction *ϕ*), of pure multivalent protein system versus protein-oligonucleotide mixture. (C) Snapshots of direct coexistence simulations of pure protein (top) and protein-oligonucleotide mixture (bottom) at 1*/E*_*prot*-*prot*_ = 0.095.

All potentials in our minimal patchy model are continuous, and can be conveniently implemented in parallelised molecular dynamics (MD) software. Accordingly, the so-called MD-Patchy model^37^ can probe condensed-matter properties in an efficient manner. We employ direct coexistence simulations^38–40^ to compute the phase diagrams of our protein-oligonucleotide mixtures; i.e. both liquid phases are simulated via MD in the same simulation box.^37^

Specifically, we measure the concentrations (packing fractions) of the protein-rich and protein-poor phases as a function of protein-protein interaction strength. The packing fraction (*ϕ*) of a molecule (*i*) in a phase is defined as the volume of the phase occupied by *i* (*ϕ* = *NV*_*i*_*/V* ∝ *C*_*i*_; where *N* is the number of molecules *i* and *C*_*i*_ is the concentration). LLPS in cells occur over narrow temperature ranges, therefore, we assess the phase behaviour of our mixtures by varying the inter-protein interaction strengths (particularly, 1*/E*_*prot*-*prot*_; the inverse interaction strength) at a fixed temperature.

For a particular protein, the compositions of the coexisting protein-rich and protein-poor phases define the range of concentrations for which LLPS takes place (Fig. 1B). Having established this reference phase diagram, we then assess the effect of adding oligonucleotides or different proteins on the location of the phase boundaries (Fig. 1C). This approach allows us to obtain general rules for how proteins and oligonucleotides partition into two-liquid phases.

Readers are referred to Ref. 37 and the Methods Section for further details on the MD-Patchy model and the simulation parameters.

### Oligonucleotides are high-valency molecules that can act as super-scaffolds for LLPS

Many proteins that undergo LLPS possess RNA-recognition motifs (RRMs).^20,26,27^ It has been suggested that LLPS is aided by multivalent interactions between such proteins and RNA. For example, Molliex and coworkers^26^ showed that, while RNA-binding protein (RBP) hnRNPA1 was able to phase-separate on its own, the LLPS of hnRNPA1 was significantly enhanced in the presence of RNA. This effect was marked by a dramatic decrease in the concentration of hnRNPA1 required for phase separation.^26^ Similar results were also reported by Lin and colleagues.^20^ RNA was also found to drive droplet formation of RBP Whi3 at physiological protein concentrations.^18,27^ In another study, Zhang et al.^27^ demonstrated that the RRM of Whi3 was required for RNA-mediated phase separation.

To decipher the molecular mechanism for oligonucleotide-driven LLPS, we investigated the effects of such polymers on the phase boundary of a single type of multivalent protein (i.e. the 3-valency protein). The protein can self-associate via three binding sites on its surface. At different inter-protein interaction strengths, we computed the phase boundaries of the pure protein system (Fig. 1B). We then calculated the coexistence curve and the minimal interaction strength needed for phase separation (i.e. the critical point). Above the critical point (i.e. if the interaction strength divided by the thermal energy is below a critical threshold) the pure protein system forms a single well-mixed phase (top panel in Fig. 1C). At larger interaction strengths, the system separates into a protein-rich and protein-poor phase.

We then added an oligonucleotide (made up of 40 hard monomeric units) and simulated the new protein-oligonucleotide mixture above the critical point of the pure system (Fig. 1B). Consistent with experimental studies of RNA-RBP mixtures, the oligonucleotide was found to promote phase separation of the protein (bottom panel in Fig. 1C). That is, a higher critical inverse interaction strength and a broader coexistence region was obtained for the mixture versus the pure system (Fig. 1B). Hence, at a given inter-protein interaction strength the saturation concentration for LLPS decreased due to the presence of the oligonucleotide.

As in the case of the Whi3-RNA mixture,^18,27^ our multivalent proteins interact with the oligonucleotide via different sites than those used for protein-protein association. Accordingly, oligonucleotides act a high-valency molecules that effectively enhances the average valency of the proteins. Indeed the average number of protein-protein bonds in the pure system is 1.00, compared to 1.13 bonds per protein molecule when the oligonucleotide is added at an inverse interaction strength of 0.09. In recent simulations, it was shown that marginal variations in average valency may lead to significant changes in phase behaviour (see Fig. 9b in Ref. 37). Li et al.^5^ also obtained a strong correlation between the phase boundary and the valency of interacting molecules for engineered SH3_m_-PRM_n_ protein mixtures. Importantly, Rao and Parker^41^ demonstrated that mRNA oligonucleotides enhanced P-body assembly by providing multiple interaction sites for certain P-body proteins. Moreover, mutations that hindered RNA-binding suppressed P-body formation.^41^ Ries et al.^42^ recently showed that polymethylated mRNAs can act as multivalent scaffolds for binding certain proteins and driving LLPS.

To further investigate the effect of oligonucleotides on the LLPS of protein systems, we defined an order parameter *Q*, which measures the number of proteins in a given protein cluster (i.e. *Q* loosely describes the size of protein condensates). A protein is considered to be in a cluster if, within a cutoff distance 4.3 Å, it has at least three neighbours. The largest protein cluster (*Q*_1_), therefore, probes the nucleation potential of the system. We first computed *Q*_1_ at different inter-protein interaction strengths for the pure protein system and in the presence of a 120-mer oligonucleotide (Fig. 2A). The critical protein-protein interaction strength (10.75 kT) for the pure protein system is indicated on the figure. Below this threshold, the largest cluster in the pure protein system contains about 52 proteins. Strikingly, there is a 10-fold increase in the size of the largest protein cluster (ca. 574 proteins) upon adding the 120-mer at extremely weak inter-protein interaction strengths (*<*10.75 kT). Additionally, when the oligonucleotide is added, we obtain a gain (of approximately 1 kT) in the stability of protein clusters. It follows that the polymer functions as an effective seed—recruiting numerous protein molecules, thereby assisting in overcoming the thermodynamic barrier for phase separation. In the literature, Falahati et al.^43^ proposed a seeding mechanism for the formation of the nucleolus. Specifically, they reported that rRNA transcription precedes nucleolus assembly,^43^ which subsequently recruits and localises nucleolar proteins– fostering protein cross-linking and eventual condensation. Our results here are consistent with a seeding-type mechanism.

**Figure 2:**
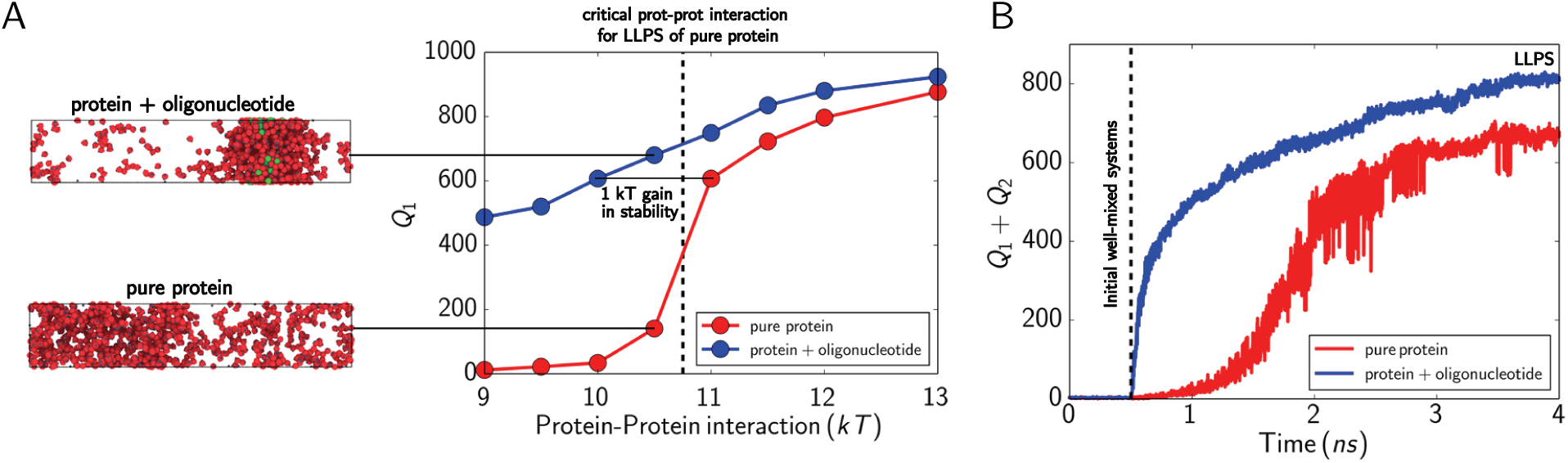
Thermodynamics and kinetics of protein cluster formation. (A) Number of proteins in largest cluster (*Q*_1_) versus protein-protein interaction strength. Snapshots of each system are included; showing a large distinct protein cluster for the protein-oligonucleotide mixture and smaller dispersed clusters in pure protein system. (B) Sum of two largest protein clusters (*Q*_1_ + *Q*_2_) as a function of time at an inter-protein bond strength of 12 kT.

Finally, we probed the evolution of the two largest protein clusters (denoted *Q*_1_ + *Q*_2_) as a function of time (Fig. 2B). In the early stages of condensation, protein clusters have a tendency to fuse and segregate often, and so the sum of the two largest clusters is a more robust condensation parameter than *Q*_1_. To obtain a homogeneous distribution of proteins in the simulation box, we initially zeroed all attractive interactions in each system. We then activated all attractive interactions; setting the protein-protein interaction strength to 12 kT. At this inter-protein bond strength, both systems undergo LLPS; therefore, *Q*_1_ + *Q*_2_ versus time directly measures the rate of condensate formation. We find that the two largest condensates form approximately four times faster in the presence of the polymer than in the pure system. Furthermore, whereas *Q*_1_ + *Q*_2_ grows in a roughly monotonic fashion in the protein-oligonucleotide mixture (blue curve in Fig. 2B), the sum fluctuates to a greater extent (red curve in Fig. 2B) in the absence of the oligonucleotide. Based on these results, we can conclude that, in addition to increasing the stability of condensates, oligonucleotides promote LLPS by reducing the kinetic barrier to condensation.

### Superscaffolds may undergo phase separation-driven compaction

We have demonstrated how oligonucleotides can modulate phase boundaries of pure protein systems. Another interesting line of inquiry pertains to the effects of phase separation on the oligonucleotides themselves. For example, how do these polymers behave within a droplet? The answers to such questions hold important implications for gene accessibility and expression. Strom et al.^36^ hypothesised that the formation of compact chromatin (heterochromatin) is driven by HP1-mediated LLPS; within HP1 phase-separated droplets chromatin is said to adopt a more compact geometry, which can contribute to transcription regulation. Two earlier studies by Nott and coworkers^12,24^ revealed that membraneless organelles preferentially recruited compact oligonucleotides over more extended/rigid ones. Moreover, in some cases rigid molecules (e.g. double-stranded DNA) were melted once inside droplets. They rationalised that shorter or compact RNA/DNA molecules were more easily incorporated into the condensate, because these molecules led to minimal disruption of the percolating network.

However, the configurational landscape of oligonucleotides inside condensed protein phases is expected to be highly diverse and to be modulated by several factors. Beyond multivalent proteins that phase-separate on their own and subsequently recruit oligonucleotides, there are proteins that do not exhibit favourable self-interactions and only phase-separate when bound to their complementary RNA/DNA strands.^18,27,44,45^ In the latter case, less compact oligonucleotides may actually favour LLPS. Hence, the nature of the proteins that form phase-separated droplets plays a critical role in influencing the conformation of constituent oligonucleotides.

Polymer chains naturally exhibit various degrees of compaction depending on their intrinsic bond stiffness (bond angle force constants; *k*_*angle*_). For example, when a 16-mer oligonucleotide is submerged in a condensed protein phase, the mean geometry of the polymer varies from extended to folded (compact) as the bond stiffness is decreased (Fig. 3A). However, the relative binding affinities of the various components in biomolecular condensates may further affect the averaged structures of sequestered oligonucleotides.

**Figure 3:**
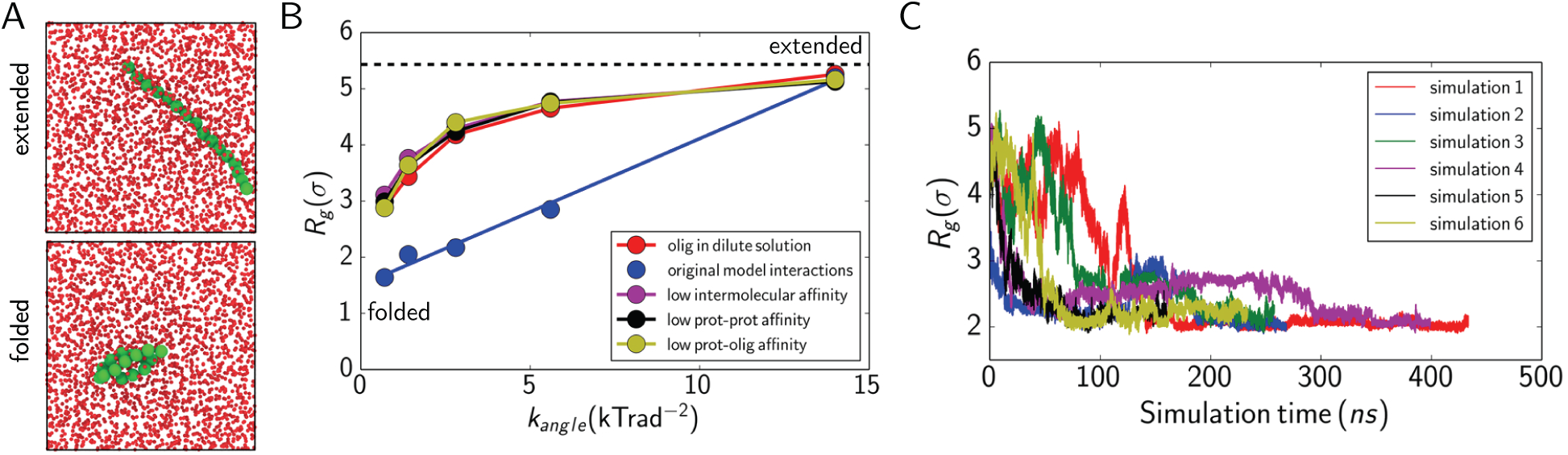
Radius of gyration (*R*_*g*_; *σ* = molecular diameter of one bead) of oligonucleotides within biomolecular condensates. (A) Snapshots of an extended (top) versus a folded/collapsed polymer (bottom) within a condensate. The radii of the proteins (red) have been reduced for clarity. (B) *R*_*g*_ as a function of angular force constant (*k*_*angle*_ in units of KT). The intermolecular interactions are modulated to investigate different biologically relevant scenarios: (red) oligonucleotide in an infinitely dilute solution (or vacuum; “control”); (blue) original model interactions (i.e. weak attractive interactions between all components); (purple) low intermolecular affinity (repulsive forces dominate); (black) low protein-protein affinity (negligible attractive protein-protein interactions); (yellow) low protein-oligonucleotide affinity (negligible attractive protein-oligonucleotide interactions). (C) *R*_*g*_ versus simulation time for six independent protein-oligonucleotide mixtures. In all simulations, the oligonucleotide becomes more compact without minimising its total interaction energy (see *SI Appendix* Fig. A.1).

Due to the simplicity of our model, we were able to investigate the impact of different biological scenarios on the behaviour of oligonucleotides within biomolecular condensates. Specifically, we measure the radius of gyration of an oligonucleotide in a condensed phase as a function of the intermolecular interactions of the phase components (Fig. 3B): (1) Original model interactions (weak attractive intermolecular interactions); (2) Low intermolecular affinity (repulsive forces dominate); (3) Low protein-protein affinity (negligible inter-protein attractions); (4) Low protein-oligonucleotide affinity (negligible protein-polymer attractions). As a control, we also simulate the oligonucleotide in an infinitely dilute solution (or in vacuum; red curve in Fig. 3B).

Interestingly, we find that the polymer is only more compact when all components interact via weak attractive interactions (blue curve in Fig. 3B). In the other cases, where some or all attractive interactions are effectively zeroed, the polymer essentially behaves as if it were suspended in a good solvent. These results demonstrate that polymer compaction in condensates is not trivial; rather, this process is likely highly cooperative—depending strongly on the precise interplay between intermolecular interactions.

We then investigated the origin of the phase separation-driven compaction observed in our model (blue curve in Fig. 3B). To this end, we ran six independent simulations of the protein-oligonucleotide mixture (each starting from a different initial configuration), and analysed the variation in the total interaction energy along each trajectory. In all simulations, the oligonucleotide equilibrated to a more compact geometry (Fig. 3C). Surprisingly, we found no net gain in the interaction energy on going from an extended polymer to a more compact one (*SI Appendix* Fig. A.1). Next, we performed a decomposition of the total interaction energy; recall that the condensate is stabilised by both inter-protein bonds and protein-oligonucleotide interactions. The energy decomposition analysis revealed that neither of these energetic terms were minimised upon oligonucleotide collapse (*SI Appendix* Table A.1). Additionally, when the oligonucleotide was extended versus collapsed, on average, the same number of protein molecules were bound to it; i.e., the effective valency of the oligonucleotide was essentially invariant upon folding.

Collectively, these results indicate that, in our model, oligonucleotide compaction within the condensate is not driven by any one of the enthalpic forces in our mixture. Moreover, since the system does not minimise its interaction energy upon oligonucleotide collapse, we therefore conclude that here phase separation-driven compaction is governed by entropy. We hypothesise that the excluded volume of the a compact polymer is minimised compared to an extended one; consequently, the translational entropy of proteins is enhanced for a compact oligonucleotide. In other words, when the oligonucleotide is compact the proteins have more freedom to rearrange in the percolating network. It would be useful to test this hypothesis further in future work.

Additionally, whereas long polymers undergo compaction, short polymers tend to aggregate (*SI Appendix* Fig. A.2). Again, this phenomena may be governed by entropy and explained by the fact that polymer aggregation creates less mechanical stress on the percolating network, and minimal distortion of the condensate is achieved. Unlike long polymers, short polymers remain straight simply due to the interplay between their persistence length (i.e. the maximum length over which a polymer behaves as a shift rod) and the high density fluctuation that they can create. Overall, our results show good agreement with the findings of Nott et al.,^24^ and suggest that the compaction of oligonucleotides within membraneless organelles may be entropically driven.

### Superscaffolds regulate composition of condensates

What physical factors determine which molecules are recruited to or excluded from biomolecular condensates is another important open question. Banani and coworkers^23^ provided an initial framework to explain compositional control of membraneless organelles. Specifically, they demonstrated how low valency molecules (termed “clients”) may be recruited to condensates by binding to “scaffolds” (molecules that can phase-separate on their own). They also highlighted that the valencies and relative concentrations of scaffolds and clients played significant roles in dictating the final droplet composition. In another study, Saha et al.^16^ found that competition among certain proteins for RNA binding sites altered P-granule-like droplet composition.

Here, we investigate how various factors modulate condensate composition in protein-oligonucleotide mixtures. In particular, we examine cooperation and competition effects in multicomponent mixtures comprising scaffolds, clients and superscaffolds. First, we considered the case for a mixture containing 66% scaffolds (3-valency proteins; Fig. 4A) and 33% clients (2-valency proteins; Fig. 4A)—i.e. an average valency of 2.64 (see Simulation details in Methods Section)—with all inter-protein interaction strengths equivalent. To determine whether superscaffolds can enhance LLPS of this mixture, we added a 40-mer oligonucleotide to the client-scaffold mixture and simulated the system at an inter-protein interaction strength of 11.5 kT; at this interaction strength, the client-scaffold mixture does not undergo LLPS on its own (*SI Appendix* Fig. A.3). Upon equilibration, no LLPS was observed (Mixture I in Table 1; Fig. 4B), and the superscaffold was almost entirely coated by scaffolds (83%), despite the binding strength of both clients and scaffolds with the polymer being equal (Table 1).

**Table 1:** We start from a mixture containing 66% (3-valency proteins) scaffolds and 33% (2-valency proteins) clients (i.e. an average valency of 2.64) that undergoes LLPS. We then add a (40-mer) polymer at an inter-protein interaction strength of 11.5 kT (i.e. an inter-protein interaction strength at which the scaffold-client mixture does not undergo LLPS), modulate the cross interactions between the mixture components, and analyse whether the polymer (superscaffold) enhances LLPS. Snapshots of observed phase behaviours are given in Fig. 4.

| Mixture | I | II | III | IV |
| --- | --- | --- | --- | --- |
| client-scaffold interaction? | yes | no | yes | yes |
| client-polymer interaction? | yes | yes | no | yes |
| scaffold-polymer interaction? | yes | yes | yes | no |
| proteins coating polymer | 83% scaffolds | 98% scaffolds | 99% scaffolds | 3% scaffolds |
| bulk valency | 2.63 | 2.62 | 2.62 | 2.68 |
| polymer enhances LLPS? | no | yes | no | yes |

**Figure 4:**
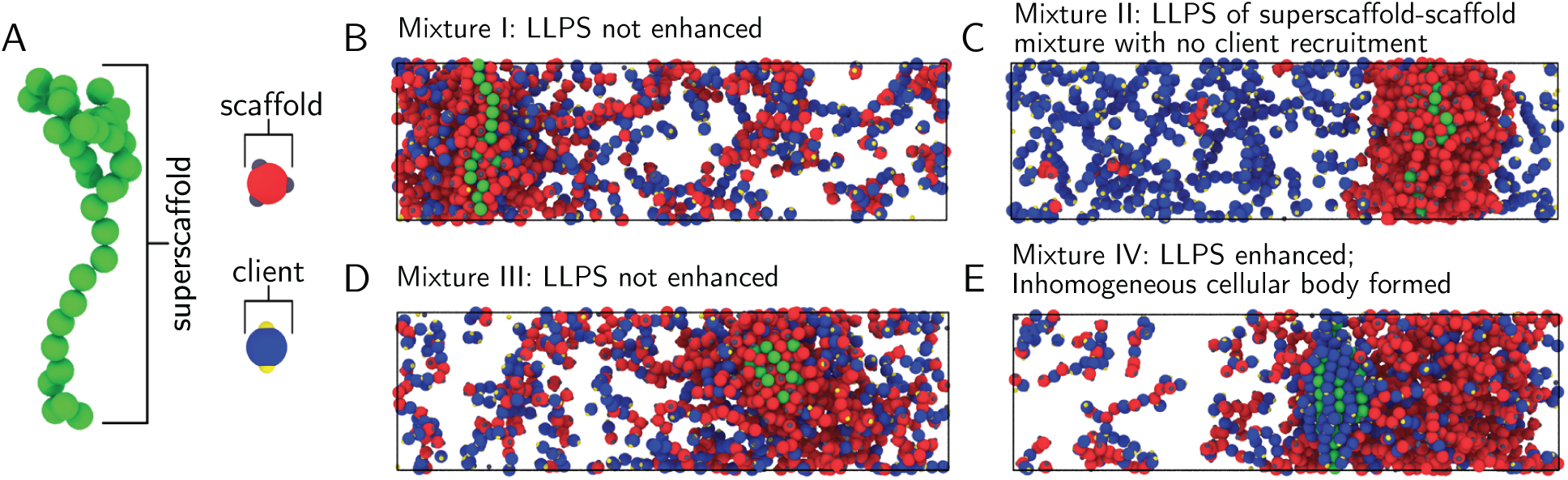
Composition of biomolecular condensates is regulated by the relative interaction strengths of components. (A) Depiction of clients (2-valency proteins), scaffolds (3-valency proteins) and superscaffold (40-mer oligonucleotide). (B)–(E) Phase behaviour of client-scaffold-superscaffold mixtures (i.e. Mixtures I–IV) described in Table 1.

Our findings can be rationalised by the fact that, despite the protein-polymer interaction strength being equal for all proteins, the dissociation constants for the higher valency proteins (which can form more bonds with surrounding molecules) are expected to be smaller; higher valency proteins therefore have a greater probability of remaining bound to superscaffolds (and bound in the percolating network, in general). Consequently, in Mixture I the scaffolds are preferentially recruited by the polymer. Once recruited, scaffolds associate with adjacent scaffolds, creating a higher concentration of clients in the remaining mixture/bulk (i.e. above the initial 33% client concentration); the average valency of the bulk phase therefore decreases (2.63). Our previous work (under review) has shown that addition of low-valency clients that are strong competitors for scaffold-scaffold interactions (like the ones in Mixture I) decreases the stability of condensates by diminishing the connectivity of the liquid-network, and that this effect is amplified as the scaffold-client ratio decreases. Since the oligonucleotide effectively decreases the scaffold-client ratio, the net effect is that droplet formation is not enhanced.

In this regard, we modelled the preceding mixture with the client-scaffold attraction switched off (Mixture II in Table 1; Fig. 4C). Despite the superscaffold being able to bind to both scaffold and clients, we observe again that it is mainly coated by scaffolds (98%). Moreover, we observe LLPS of the superscaffold-scaffold mixture, and exclusion of the clients to the diluted liquid. We hypothesised that superscaffolds preferentially recruit higher valency proteins to condensates. To test this hypothesis, a mixture containing the superscaffold and equal amounts of 4-, 3-, and 2-valency proteins was prepared. Although, the binding strength of each protein with the polymer was the same, at equilibrium the polymer was predominantly coated by the 4- and 3-valency proteins, and the droplet concentration of the proteins decreased in the order 4-*>*3-*>*2-valency protein (*SI Appendix* Fig. A.4). These results parallel experimental findings, where a strong positive correlation was found between the recruitment of FUS proteins to hydrogels and the number of tyrosine motifs in the N-terminal portion of the proteins.^46^

Next, we tested the phase behaviour of the mixture when the interaction strength between the clients and the superscaffold is effectively set to zero (Mixture III in Table 1). In this mixture (Fig. 4D), clients compete with the superscaffold for binding sites on scaffold proteins. As in Fig. 4B, no phase separation is observed. In Mixture III, 99% of the molecules coating the polymer are scaffolds; the interaction of scaffolds with the polymer and adjacent scaffolds effectively decreases the number of scaffold molecules for recruiting clients. Thus, droplet formation is not enhanced because the ratio of available scaffolds to low valency proteins (clients) that compete strongly for scaffold-scaffold interactions decreases; this is quantified by the average bulk valency decreasing to 2.62.

Finally, we switched off the attractive interaction between the scaffolds and the polymer, and assessed the effects on the phase boundaries (Mixture IV in Table 1); LLPS was observed for that mixture (Fig. 4E). Since the polymer was engineered for binding only clients, both the superscaffold and scaffolds actively recruited the lower valency proteins, and so LLPS was enhanced. Accordingly, droplet formation is promoted when superscaffolds show a preference for clients; in the final mixture, clients constitute 97% of the molecules coating the 40-mer. Ergo, the bulk valency increases (2.68) due to an excess of free protein-binding sites on scaffolds, which in turn foster recruitment of clients to droplets. In fact, Banani and coworkers^23^ noted that clients were strongly recruited to the condensate when there was a high concentration of free binding sites on cognate scaffold molecules.

In Mixture IV, we also observed a marked spacial segregation of the superscaffold and associated clients in the condensate (Fig. 4E); in contrast to the scaffold-superscaffold mixture, where the polymer was embedded within the liquid drop (bottom panel in Fig. 1C), the client-coated superscaffold was located near the droplet interface. Even in the other mixtures (Fig. 4B and 4D), where no marked protein phase separation was observed, the superscaffold was, in general, uniformly coated by the surrounding proteins. Hence, we find that spacial segregation of species within condensates can be achieved in multicomponent mixtures, and is controlled by fine-tuning the bulk valency and interaction strengths among species. Such multiphase immiscibility has been reported for the nucleolus^21^ and stress granules,^47^ where internal structuring into various sub-compartments (e.g. a core-shell) may arise. To our knowledge, the model presented here is the first to capture structural heterogeneity within droplets. Hence, this model may be useful in advancing our understanding of LLPS in multiphase biomolecular condensates.

## Conclusions

In this work, we have introduced a minimalist coarse-grained model for probing LLPS in protein-oligonucleotide mixtures. Specifically, we have studied: (*i*) the effects of nucleic acids on phase separation of proteins, (*ii*) the compaction of oligonucleotides within condensates and the possible origins of such compaction, and (*iii*) compositional control in phase-separated protein-oligonucleotide mixtures. From our simulations, fundamental rules/features relating to LLPS in these systems have emerged: (1) oligonucleotides can act as high valency molecules (“superscaffolds”) that promote phase separation by increasing the effective valency of scaffolds; (2) oligonucleotides increase the stability of condensates and decrease the kinetic barrier for nucleation; (3) oligonucleotides can undergo phase separation-driven compaction and this compaction may be entropically driven—polymer condensation maximises the translational entropy of the percolating protein network; (4) for a mixture of proteins with similar protein-superscaffold binding strengths, superscaffolds preferentially recruit higher valency proteins to condensates; (5) in a superscaffold-scaffold mixture, LLPS is hindered by clients that compete for superscaffold/scaffold binding sites. (6) spatial segregation of components within phase-separated droplets is controlled by fine-tuning the bulk valency of the mixture and interaction strengths of components.

These results improve our current understanding oligonucleotide-driven LLPS, and can be directly tested via higher resolution simulations and used to guide in vitro experiments. We hope to extend the current model to include protein flexibility effects, which may be useful in elucidating LLPS of intrinsically disordered proteins. Additionally, the model could be improved to capture sequence-dependence of protein/oligonucleotide LLPS. Nonetheless, the findings reported here represent a critical step towards a unified model for intracellular liquid-liquid phase separation.

## Methods

### MD-Patchy and polymer model

We model proteins as patchy particles by using the recently proposed MD-Patchy model.^37^ In this model, the patchy particles are (almost) hard-spheres (HS) of diameter *σ* decorated with *M* attractive sites (“patches”) on their surface. Each patch represents a particle-particle binding site. For modelling the pseudo-hard-spheres (PHS), we use a potential (*v*_*PHS*_) of the form proposed in Ref. 48:

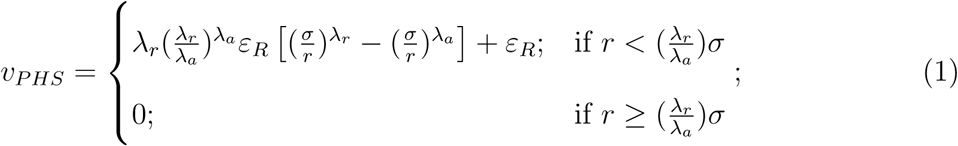

where *λ*_*a*_ =49 and *λ*_*r*_=50 are the exponents of the attractive and repulsive terms respectively, chosen for computational convenience. The interaction strength *ε*_*R*_ accounts for the energy of the pseudo hard-sphere interaction and *σ* is our unit of length. *r* denotes the centre-to-centre distance between PHS particles. For the associative sites or patches, we use a continuous attractive square-well (CSW) interaction proposed in Ref. 49:

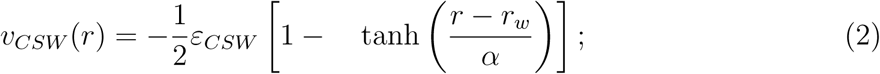

where *r* is the distance between the centres of two attractive patches, *r*_*w*_ is the radius of the attractive well, and *α* controls the steepness of the well. *ε*_*CSW*_ is the unit of energy in our simulations. To ensure that the valency of each individual patch is one (i.e. each attractive site can interact with at most one other patch at a time), we define *r*_*w*_=0.12 *σ* and *α*=0.005 *σ*.

Every colloidal patchy particle is characterised by *M* + 1 interaction sites: one central site accounting for the pseudo-hard-sphere-like interaction and *M* sites located representing the attractive sites. For our patchy particles, both the hard-sphere plus the attractive sites are defined as a multi-centre rigid body.

For modelling the polymers (oligonucleotides) we use different amounts of beads (PHS particles) joint by flexible angles and bonds. We use standard bond/angular harmonic potentials (*v*_*bonds*_ = *K*_*b*_ (*r* − *r*_0_)^2^ and *v*_*ang*_ = *K*_*a*_ (*θ* − *θ*_0_)^2^) with *K*_*b*_=100 kcal/(mol Å^2^) and *r*_0_=4.0 Å for the bonds, and *K*_*a*_=0.8 kcal/(mol *radian*^2^) and *θ*_0_=180 degrees for the angles. The value of *K*_*a*_ guarantees a high flexibility of the polymer. The interaction between the different beads of the polymer is PHS-like, whereas a standard Lennard-Jones interaction represents the interaction between the polymer beads and the patchy particles:

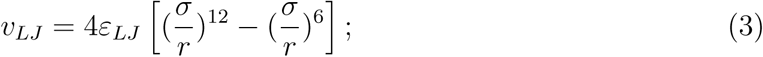

where *ε*_*LJ*_ accounts for the depth of the attractive interaction (*ε*_*LJ*_ =1.07 kcal/mol), *r* is the distance between the HS particles of the patchy particles and the beads of the polymer, and *σ* is the molecular diameter of the beads, which is the same for patchy particles and beads, *σ*=3.405 Å. We set this value of *ε*_*LJ*_ because this is the minimum energy needed to ensure a stable attractive affinity between the polymer and the patchy particles. However, with reasonably higher values of *ε*_*LJ*_ similar results are obtained.

Although the choice of the mass is irrelevant for equilibrium simulations, we set the mass of the PHS particles (polymer beads and central particle of the patchy particles) as 20 amu, while the associative sites of the patchy particles are assigned a mass of 1 amu.

We use the following reduced units for our results: 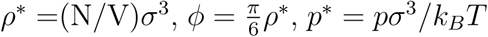 and time as 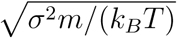 (where *T* = 119.81*K*). In order to keep the isotropic HS-like interaction in our simulations as similar as possible to a pure HS interaction, we follow Ref. 37 and fix *k*_*B*_*T/ε*_*R*_ at a value 1.5.^38,48^ We then control the effective strength of the associating sites by varying *ε*_*CSW*_, such the reduced temperature is defined as *T* * = *k*_*B*_*T/ε*_*CSW*_. At *k*_*B*_*T/ε*_*R*_=1.5 (i.e. *T* = 179.71*K*), the attractive interaction strength between the polymer beads and the patchy particles, *ε*_*LJ*_, is about 3*k*_*B*_*T*.

### Simulation details

Given that all employed potentials are continuous and differentiable, all our simulations were performed in LAMMPS.^50^ The numerical details are as follows: the timestep chosen for the Verlet integration of the equations of motion was Δ*t*=0.0004 in reduced units (that corresponds to 0.5 fs in the LAMMPS input file). The cut-off radius for both HS and CSW interactions is 1.175 *σ*. We use the Nosé-Hoover thermostat^51,52^ for NVT simulations with a typical relaxation time of 0.125 in reduced units. For NPT simulations, the Nosé-Hoover barostat was used^53^ (in combination with the Nosé-Hoover thermostat) with a relaxation time of 0.45 in reduced units. To account for the rotational motion of the patchy particles, we described the colloidal particles as rigid bodies, using the method implemented in LAMMPS,^54^ which was also employed in Ref. 37 and recent work (under review).

To evaluate phase diagrams, we use Direct Coexistence (DC) simulations.^38–40^ DC simulates coexistence by preparing periodically extended slabs of the two coexisting phases, e.g. the liquid and the vapour, in the same simulation box. We prepare the initial configurations by using the same procedure described in Ref. 37. When introducing the polymer, we add it to the vapour phase to avoid particle overlap between polymer and patchy particles.

For evaluating the phase diagram of *M* = 3 (i.e. 3-valency proteins) with the polymer, we use systems containing 1000 patchy particles. For the binary mixtures of *< M >*= 2.64 (where *< M >*= (*N*_*particles*1_*M*_*sites*1_ + *N*_*particles*2_*M*_*sites*2_)*/N*_*total*_), we used 768 and 432 patchy particles for *M* = 3 and *M* = 2, respectively, and a polymer chain of 40 monomers (beads). For the ternary mixture of *M* = 4, *M* = 3 and *M* = 2, 400 patchy particles of each type were used, along with a 40 monomer polymer. To compute the radius of gyration, a mixture of 10000 patchy particles and a polymer of 16 monomers were used. Unlike the other cases, here a cubic simulation box was used to avoid self-interactions of the polymer through the periodic boundaries.

## Acknowledgement

This project was funded by the European Research Council (ERC) under the European Union Horizon 2020 research and innovation programme (grant agreement No 803326). R.C.-G. is an Advanced Research Fellow from the Winton Programme for the Physics of Sustainability. J.A.J is a King’s College Research Fellow. J.R.E. is an Oppenheimer Research Fellow and an Emmanuel College Research Fellow. A.G. is funded by an EPSRC studentship (EP/N509620/1). This work was performed using resources provided by the Cambridge Tier-2 system operated by the University of Cambridge Research Computing Service (http://www.hpc.cam.ac.uk) funded by EPSRC Tier-2 capital grant EP/P020259/1.

## Supporting Information Appendix

**Figure A.1:**
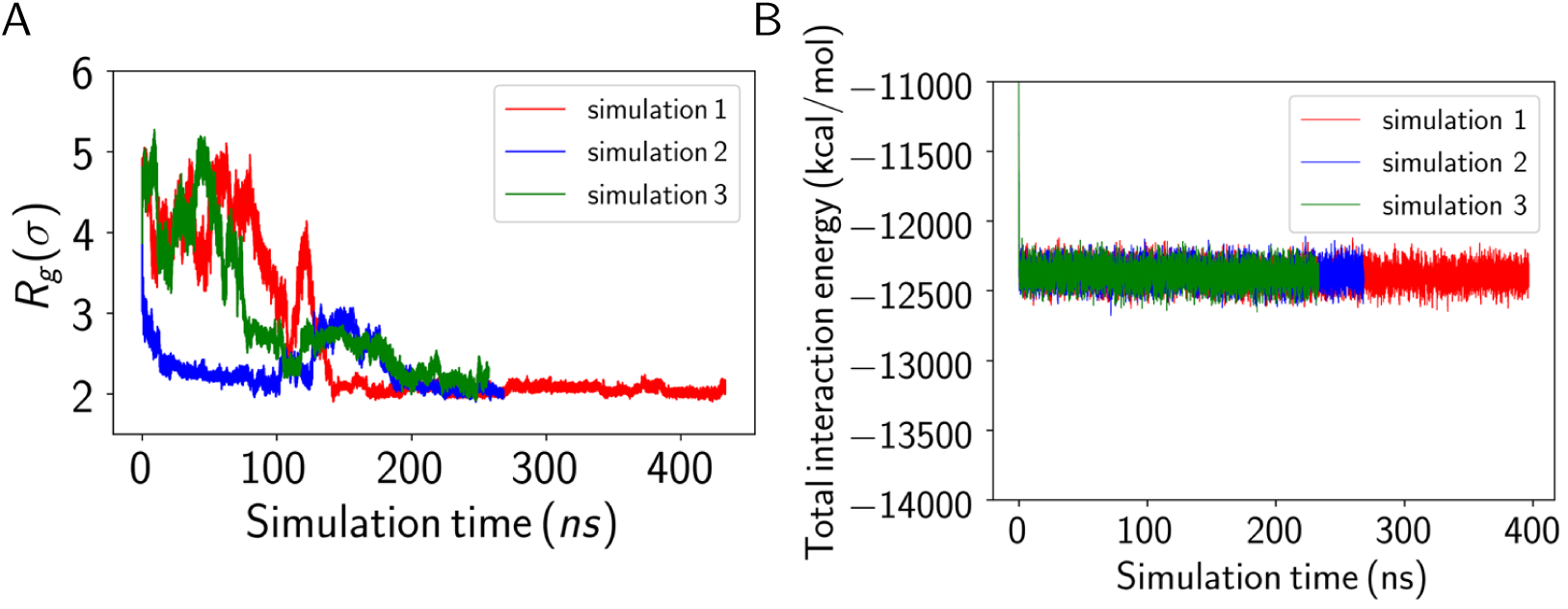
Variation in (A) radius of gyration (*R*_*g*_) and (B) total interaction energy as a function of simulation time for three independent simulations composed of 3-valency proteins (10000) and a 16-mer oligonucleotide. In all cases, the polymer converged to a more compact geometry. However, there was no net gain in the total interaction strength on going from the extended polymer to a more compact one.

**Table A.1:** Energy decomposition for a mixture of 3-valency proteins (10000) and a 16-mer oligonucleotides. All energies are given in kcal/mol (Key: Total potential energy = *E*_*PE*_; Protein-protein interaction strength = *E*_*prot*-*prot*_; Number of protein-oligonucleotide interactions = no. prot-olig; Protein-oligonucleotide interaction strength = *E*_*prot*-*olig*_; Bond angle energy = *E*_*angle*_). Here, we monitor the variation of the total interaction strength (and corresponding potential energy terms) in the earlier and latter parts of the simulation (i.e. before and after compaction of the polymer).

| $R_g(\text{\AA})$ | $E_{PE}$ | $E_{prot-prot}$ | no. prot-olig | $E_{prot-olig}$ | $E_{angle}$ |
| --- | --- | --- | --- | --- | --- |
| 14.65 | -12363.9 | -12370.9 | 70.6 | -168.35 | 5.2 |
| 7.02 | -12355.9 | -12352.0 | 80.6 | -184.05 | 8.1 |

**Figure A.2:**
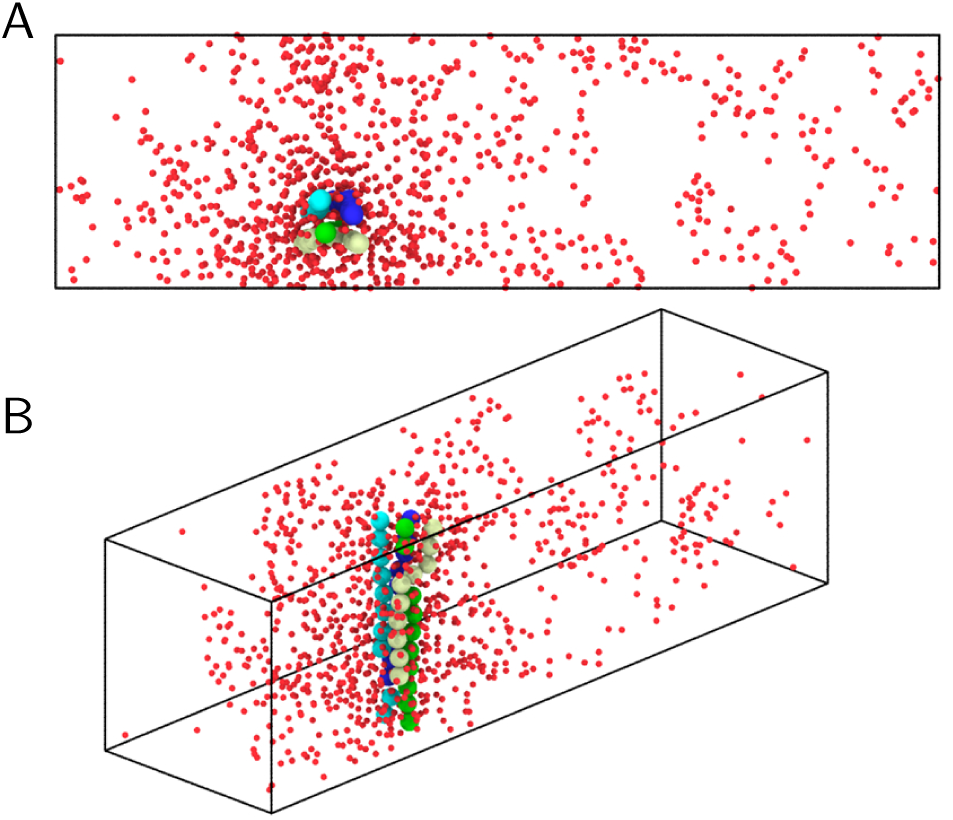
Snapshots of direct coexistence simulation of a mixture of 1000 3-valency proteins (radii reduced for clarity) and four 10-mer oligonucleotides. Although the cross interaction between the polymers is repulsive, the polymers aggregate; this arrangement creates less mechanical stress on the percolating network, and minimal distortion of the condensate. Unlike long polymers, short polymers remain straight due to the interplay between their persistence length and the high density fluctuation that they can create. Two different views are given: (A) Top and (B) Perspective.

**Figure A.3:**
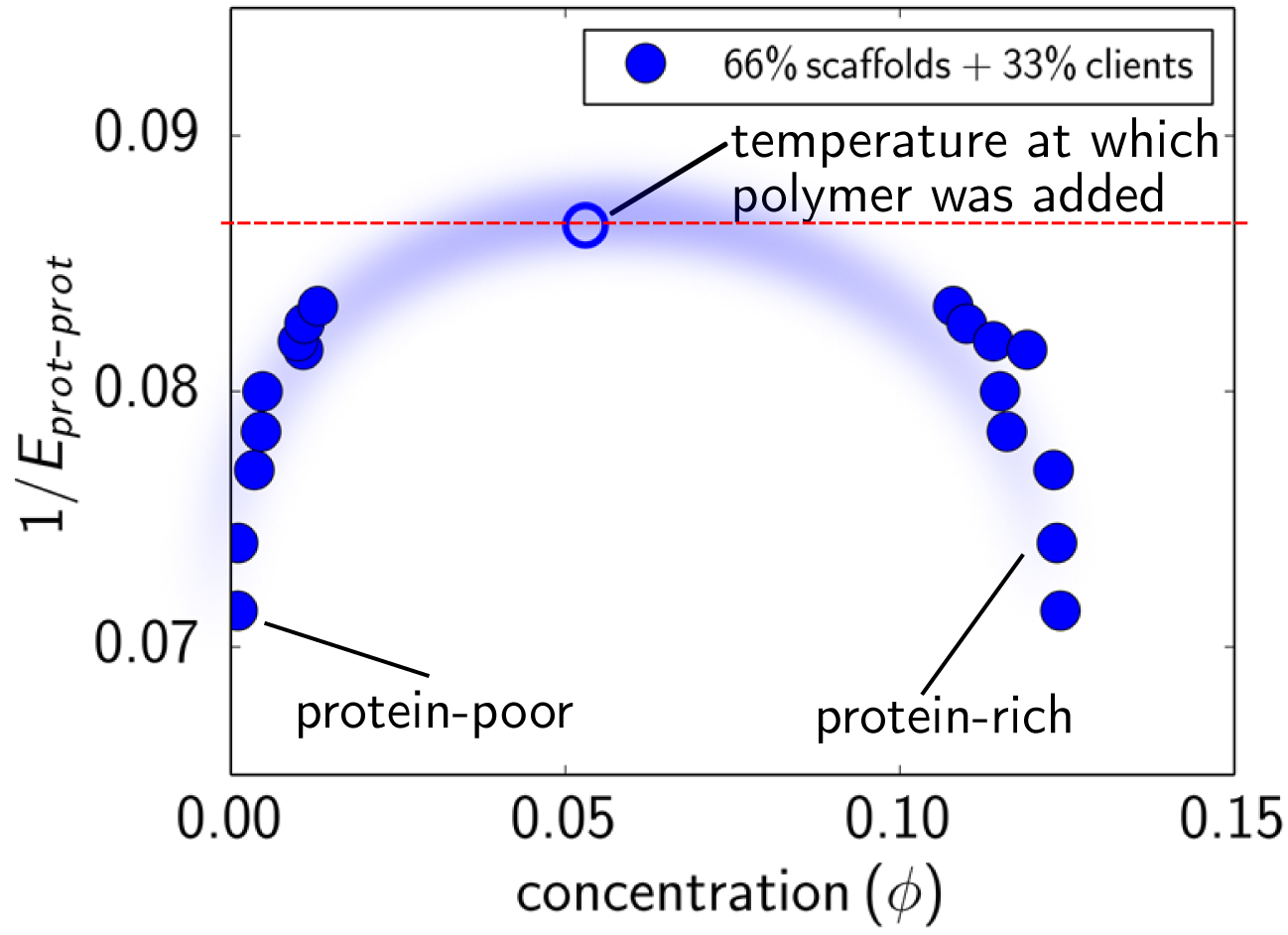
Phase diagram for a mixture containing 66% (3-valency proteins) scaffolds and 33% (2-valency proteins) clients. The inter-protein interaction strength (or temperature) at which the polymer was added for mixtures I–IV (described in the main paper) is indicated on the plot.

**Figure A.4:**
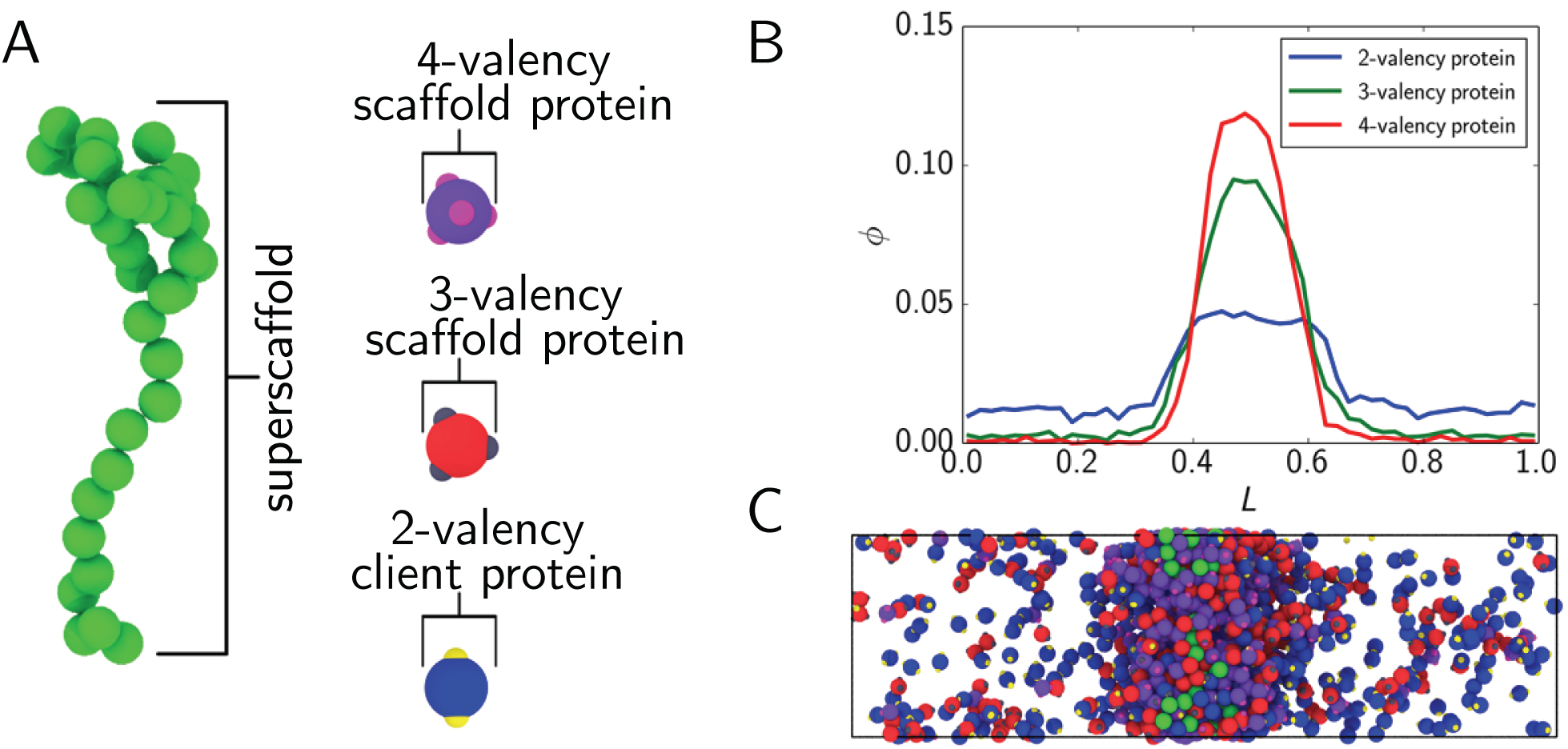
Superscaffolds preferentially recruit higher valency proteins to condensates. (A) Minimalist coarse-grained model used to simulate interactions between proteins (scaffolds and clients) and oligonucleotide (superscaffold). (B) Protein concentration (i.e. packing fraction *ϕ*) as a function of the simulation box length (*L*) for protein components in a mixture containing 1/3 each 4-,3-, and 2-valency proteins and a 40-mer oligonucleotide. (C) Snapshot of direct coexistence simulation of mixture in (B).

